# Theory for transitions between exponential and stationary phases: universal laws for lag time

**DOI:** 10.1101/135665

**Authors:** Yusuke Himeoka, Kunihiko Kaneko

## Abstract

The quantitative characterization of bacterial growth has attracted substantial research attention since Monod’s pioneering study. Theoretical and experimental work have uncovered several laws for describing the exponential growth phase, in which the number of cells grows exponentially. However, microorganism growth also exhibits lag, stationary, and death phases under starvation conditions, in which cell growth is highly suppressed, for which quantitative laws or theories are markedly underdeveloped. In fact, the models commonly adopted for the exponential phase that consist of autocatalytic chemical components, including ribosomes, can only show exponential growth or decay in a population, and thus phases that halt growth are not realized. Here, we propose a simple, coarse-grained cell model that includes an extra class of macromolecular components in addition to the autocatalytic active components that facilitate cellular growth. These extra components form a complex with the active components to inhibit the catalytic process. Depending on the nutrient condition, the model exhibits the typical transitions among the lag, exponential, stationary, and death phases. Furthermore, the lag time needed for growth recovery after starvation follows the square root of the starvation time and is inversely related to the maximal growth rate. This is in agreement with experimental observations, in which the length of time of cell starvation is memorized in the slow accumulation of molecules. Moreover, the lag time distributed among cells is skewed with a long time tail. If the starvation time is longer, an exponential tail appears, which is also consistent with experimental data. Our theory further predicts a strong dependence of lag time on the speed of substrate depletion, which can be tested experimentally. The present model and theoretical analysis provide universal growth laws beyond the exponential phase, offering insight into how cells halt growth without entering the death phase.

## Introduction

Quantitative characterization of a cellular state, in terms of the cellular growth rate, concentration of external resources, as well as abundances of specific components, has long been one of the major topics in cell biology, ever since the pioneering study by Monod [1]. Such studies have been developed mainly by focusing on the microbial exponentially growing phase, in which the number of cells grows exponentially (this phase is often termed the *log phase* in cell biology, but considering the focus on exponential growth, we here adopt the term ”exponential phase” throughout). This work has uncovered somewhat universal growth laws, including Pirt’s equation for yield and growth [2] and the relationship between the fraction of ribosomal abundance and growth rate (experimentally demonstrated by Schaechter *et al.*[3], and theoretically rationalized by Scott *et al.* [4]), among others [5–8], in which the constraint to maintain steady growth leads to general relationships[9–11].

In spite of the importance of the discovery of these universal laws, cells under poor conditions exhibit different growth phases in which such relationships are violated. Indeed, in addition to the death phase, cells undergo a stationary phase under conditions of resource limitation, in which growth is drastically suppressed. Once cells enter the stationary phase, a certain time span is generally required to recover growth after resources are supplied, which is known as the lag time. There have been extensive studies conducted to characterize the stationary phase, including the length of lag time for resurrection and the tolerance time for starvation or antibiotics [12–14], and specific possible mechanisms for phase transitions have been proposed [15–17]. Furthermore, recent experiments have uncovered the quantitative relationships of lag time and its cell-to-cell variances[18, 19]. For example, the lag time was shown to depend on the length of time the cells are starved. This implies that the stationary phase is not actually completely stationary but that some slow changes still progress during the starvation time, in which cells “memorize” the starvation time. Hence, a theory to explain such slow dynamics is needed that can also characterize the phase changes and help to establish corresponding quantitative laws.

The existence of these phases and lag time are ubiquitous in bacteria (as well as most microorganisms). Hence, we aimed to develop a general model that is as simple as possible, without resorting to detailed specific mechanisms, but can nonetheless capture the changes among the lag, exponential, stationary, and death phases. We first describe a simple model for a growing cell, which consists of an autocatalytic process driven by active chemical components such as ribosomes. However, this type of model with autocatalytic growth from substrates and their derivatives that is adopted for the exponential phase is not sufficient to represent all phases, as the autocatalytic process either grows exponentially or decays toward death, and thus does not account for a halting state with suppressed growth corresponding to the stationary phase. Therefore, to go one step further beyond the simplest model, we then consider the addition of an extra class of components that do not contribute to catalytic growth. Still, even the inclusion of this extra class of components cannot fully account for the transition to the stationary phase. Therefore, we further considered the interaction between the two classes of components. Here, we propose a model that includes the formation of a complex between these two types of components, which inhibits the autocatalytic process by the active components. We show that the model exhibits the transition to the stationary phase with growth suppression. By analyzing the dynamics of the model, we then uncover the quantitative characteristics of each of these phases in line with experimental observations, including the bacterial growth curve, quantitative relationships of lag time with starvation time and the maximal growth rate, and the exponentially tailed distribution of lag time. The proposed model also allows us to derive several experimentally testable predictions, including the dependence of lag time on the speed of the starvation process.

### Model

Since molecules that contribute to autocatalytic processes are necessary for the replication of cells, models for growing cells generally consist at least of substrates (*S*) and active components (noted as “component A” hereafter) that catalyze their own synthesis as well as that of other components. For example, in the models developed by Scott *et al.*[4] and Maitra *et al.* [20], component A corresponds to ribosomes, whereas several models involving catalytic proteins have also been proposed[10, 21– 24]. This class of models provides a good description of the exponential growth of a cell under the condition of sufficient substrates availability; however, once the degradation rate of component A exceeds its rate of synthesis under a limited substrate supply, the cell’s volume will shrink, leading to cell death. Hence, a cell population either grows exponentially or dies out, and in this cellular state it is not possible to maintain the population without growth. However, cells often exhibit suppressed growth under substrate-poor conditions, even at a single-cell level [12, 13, 18], as observed in the stationary phase. Such cells that neither grow exponentially nor go toward death cannot be modeled with cell models that only consider autocatalytic processes[4, 10, 20–24].

Therefore, to model a state with such suppressed growth, it is important to consider additional chemical species, i.e., macromolecules that do not contribute to autocatalytic growth, in addition to the substrates (*S*) and component A (*A*) that are commonly adopted in models of cell growth. Component A represents molecules that catalyze their own growth such as ribosomes, and can include metabolic enzymes, transporters, and growth-facilitating factors. Component B represents waste products or can be other molecules that are produced with the aid of component A but do not facilitate growth. Thus, the next simplest model is given by

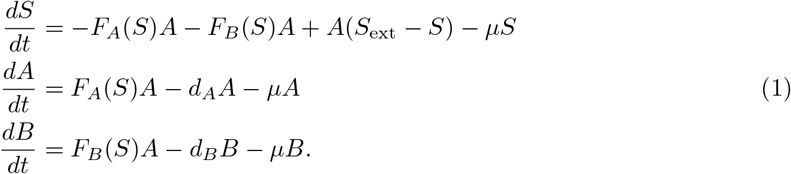

Here, *S*_ext_ and *S* indicate the concentrations of the extracellular and intracellular substrate, respectively. The concentration of the intracellular substrate determines the synthesis rate of the active and non-autocatalytic proteins *F_A_* and *F_B_*, respectively. All chemical components are diluted due to the volume growth of a cell.

In addition to dilution, macromolecules (A and B) are spontaneously degraded with slow rates (*d_A_ and *d_B_*). In this model, the cell takes up substrates from the external environment from which component A and the non-growth-facilitating component B are synthesized. These syntheses, *S*_ext_ ↔ *S,S* → *A*, and *S* → *B*, as well as the uptake of substrates take place with the aid of catalysis by component A. Then, by assuming that the synthesized components are used for growth in a sufficiently rapid period, the growth rate is set to be proportional to the synthesis rate of component A. Hence, the dilution rate *μ* of each component due to cell volume growth is set as *μ* = *F_A_A**.

Now, if the ratio *F_A_*/*F_B_* does not depend on the substrate concentration *S*, the fraction *A*/*B* also does not depend on *S*, and the model is reduced to the original autocatalytic model; thus, the phase change to suppressed growth is not expected. Then, by introducing the *S*-dependence of *F_A_*/*F_B_* to reduce the rate of component A with the decrease in the substrate condition, we first tested whether the transition to a suppressed growth state, as in the stationary phase, occurs under a substrate-poor condition, by setting *F_A_*/*F_B_* to decrease in proportion to the change in *S* (i.e., 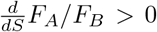). However, in this case, it is straightforwardly confirmed that there is no transition to a suppressed growth state. That is, the cells always grow exponentially without any slowing-down process, as the decrease in *S* simply influences the growth rate *μ*, while the presence of *B* does not influence the dynamics of *A*. (see also Appendix A).

Thus, we need to introduce an interaction between component A and the non-growth-facilitating component B. Although complicated interactions that may involve other components could be considered, the simplest and most basic interaction that can also provide a basis for considering more complex processes would be formation of a complex between A and B given by the reaction *A* + *B* ↔ *C*. This results in inhibition of the autocatalytic reaction for cell growth, as complex C does not contribute to the activity for the autocatalytic process. A schematic representation of the present model is shown in Fig. 1(a). Thus, our model is given by

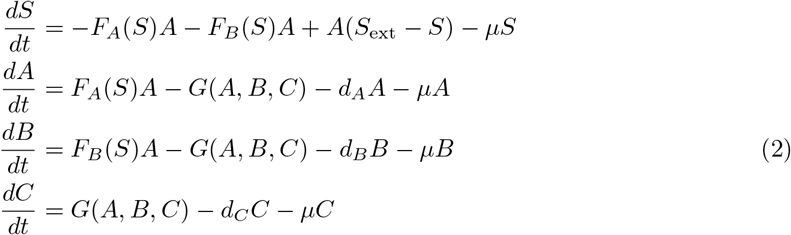

where *G*(*A, B, C*) denotes the reaction of complex formation, given by *k_p_AB* − *k_m_C*. The catalytic activity of component A is inactivated due to the formation of complex C. Here, the complex has higher stability than that of other proteins (*d_C_* is smaller than *d_A_* and *d_B_*)[25] From Eq. (2), by summing up 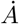 and *Ċ*, we obtain 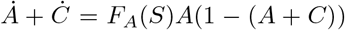 if *d_A_* and *d_C_* are zero (or negligible). This means that once the cell reaches any steady state, the relationship *A* + *C* = 1 is satisfied as long as *A* and *F_A_*(*S*) are not zero. We use this relationship and eliminate *C* by substituting *C* = 1 − *A* for the following analysis.

**FIG 1.**
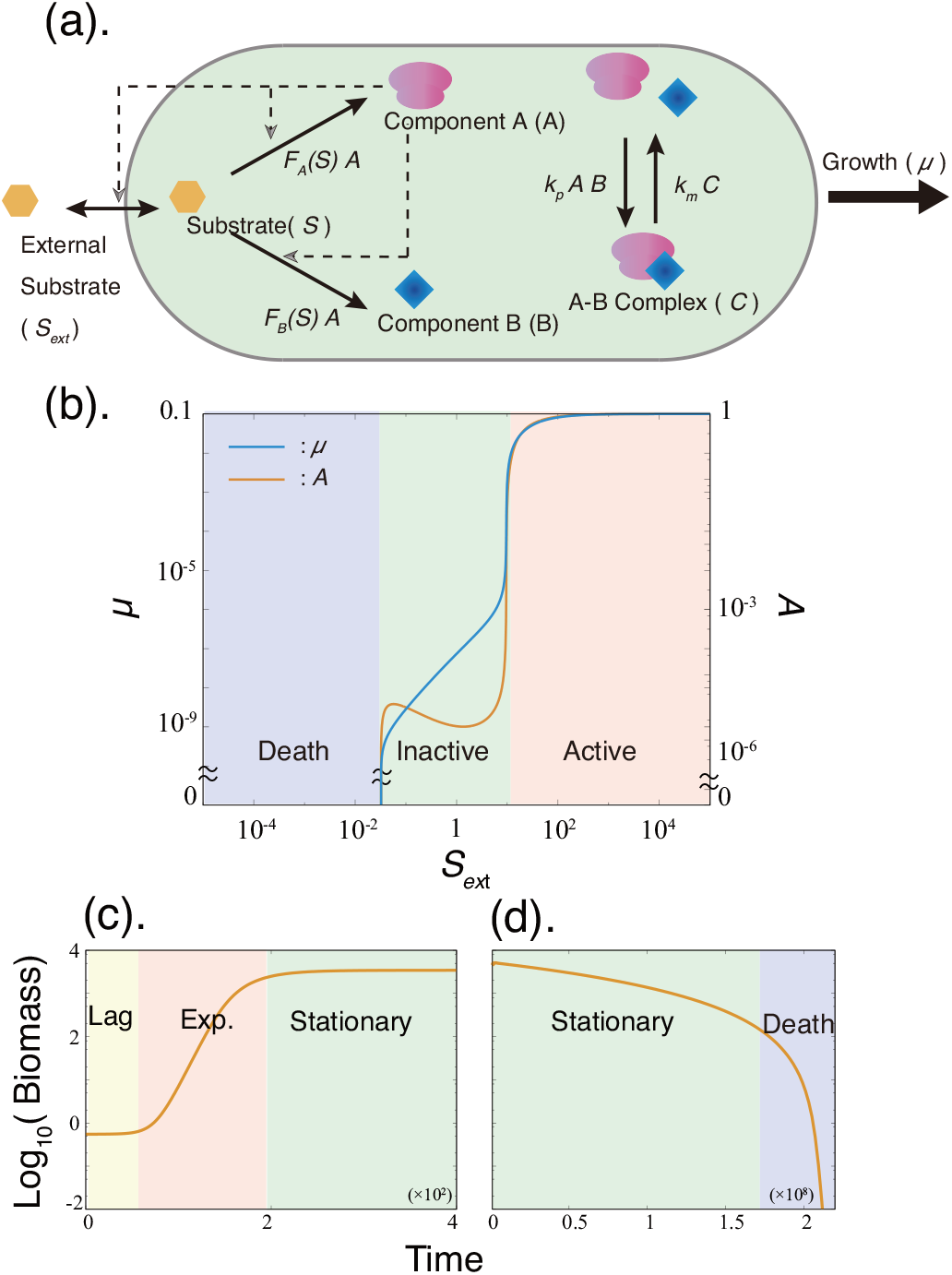
(a) Schematic representation of the components and reactions in the present model. The concentration of each chemical changes according to the listed reactions. In addition, chemicals are spontaneously degraded at a low rate, and become diluted due to volume expansion of the cell. (b) Steady growth rate and the concentration of component A are plotted as functions of the external concentration of the substrate. (c and d) Growth curve of the model. Parameters were set as follows: *v* = 0.1, *k_p_* = 1.0, *k_m_* = 10^−6^, *K* = 1.0, *K_t_* = 10.0, *d_R_* = *d_B_* = 10^−5^, *d_C_* = 10^−12^. The detailed numerical method for (c) and (d) is given in Appendix C.

One plausible and straightforward interpretation of *B* is misfolded or mistranslated proteins that are produced erroneously during the replication of component A. Such waste molecules often aggregate with other molecules[26–28]. Alternatively, B components can be specific molecules such as HPF and YfiA[29–31], which inhibit catalytic activity by reacting with component A.

With regards to the formation of error or “waste” proteins, there are generally intracellular processes for reducing their fraction. These include kinetic proofreading, molecular chaperones, and protease systems. These error-correction or maintenance systems are energy-demanding, and require the non-equilibrium flow of substrates[32, 33]. Therefore, the performance of these mechanisms is inevitably reduced in a substrate (energy source)-poor environment. Thus, it naturally follows that the ratio of the synthesis of active proteins to wastes is an increasing function of the substrate concentration, i.e., 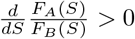. In the present model, we assume that this ratio increases with the concentration and becomes saturated at higher concentrations, as in Michaelis-Menenten’s form, and choose 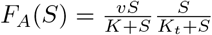 and 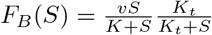, for example.

Note that almost all the results to be presented in this manuscript are obtained as long as *F_A_* ≫ *F_B_* holds for the nutrient-rich condition and *F_A_* ≪ *F_B_* for the nutrient-poor condition (see the section ”Remarks on the choice of parameters to fit the experimental data” and *Supplemental Information*). Under this condition, specific choice of the form of *F_A_* and *F_B_* is not important.

This *S*-dependence of *F_A_*/*F_B_* would be biologically plausible both for the interpretations of component B as specific inhibitory proteins or ”waste” (mistranslation) proteins. For the first interpretation, it is reported that such proteins related with the stationary phase (HPF, YfiA and others) are induced under stress condition such as starvation[29, 30, 34, 35], and thus it is suggested that *F_A_* ≫ *F_B_* (*F_A_* ≪ *F_B_*) for a large (small) amount of *S*, repectively. On the other hand, by adopting the latter, waste, interpretation, *F_A_*(*S*) and *F_B_*(*S*) close to the above Michaelis-Menten’s form is derived, by considering a proofreading mechanism to reduce the mistranslation (see also Appendix B).

Here we also note although the *S*-dependence of *F_A_*/*F_B_* is relevant to derive quantitative laws on the lag-time in agreement with experimental observation, it is not required just to show a transition to a suppressed growth state, as briefly discussed later (see Discussion).

## RESULTS

### Growth phases

The steady state of the present model exhibits three distinct phases as a function of the external substrate concentration *S*_ext_ (Fig. 1(b)), as computed by its steady-state solution. The three phases are distinguished by both the steady growth rate and the concentration of component A, which are termed as the active, inactive, and death phases, as shown in Fig.1, whereas the growth rate shows a steep jump at the boundaries of the phases. The phases are characterized as follows. **(i)** In the active phase, the highest growth rate is achieved, where there is an abundance of component A molecules, which work freely as catalysts. **(ii)** In the inactive phase, the growth rate is not exactly zero but is drastically reduced by several orders of magnitude compared with that in the active phase. Here, almost all of the component A molecules are arrested through complex formation with component B, and their catalytic activity is inhibited. **(iii)** At the death phase, a cell cannot grow, and all of components A, B, and complexes go to zero. In this case, the cell goes beyond the so-called “point of no return” and can never grow again, regardless of the amount of increase in *S*_ext_, since the catalysts are absent in any form. (As will be shown below, the active and inactive phases correspond to the classic exponential and stationary phases; however, to emphasize the single-cell growth mode, we adopt these former terms for now).

The transition from the active to inactive phase is caused by the interaction between components A and B. In the substrate-poor condition, the amount of component B exceeds the total amount of catalytic proteins (*A* + *C*), and any free component A remaining vanishes. Below the transition point from the inactive to death phase, the spontaneous degradation rate surpasses the synthesis rate, at which point all of the components decrease. This transition point is simply determined by the balance condition *F_A_* = *d_A_*. Hence, if *d_A_* is set to zero, the inactive-death transition does not occur.

We now consider the time series of biomass (the total amount of macromolecules) that is almost proportional to the total cell number, under a condition with a given finite resource, which allows for direct comparison with experimental data obtained in a batch culture condition (Fig. 1(c and d)). To compute the time series of biomass, we used a model including the dynamics of *S*_ext_ in addition to *S, A, B*, and *C*. Details of this model are shown in Appendix C. In the numerical simulation, the condition with a given, finite amount of substrates corresponding to the increase of cell number is implemented by introducing the dynamics of the external substrate concentration into the original model. Here, *S*_ext_ is decreased as the substrates are replaced by the biomass, resulting in cell growth. At the beginning of the simulation, the amount of biomass (i.e., cell number) stays almost constant, and then gradually starts to increase exponentially. After the phase of exponential growth, the substrates are consumed, and the biomass increase stops. Then, over a long time span, the biomass stays at a nearly constant value until it begins to slowly decrease. Finally, the degradation dominates and the biomass (cell number) falls off dramatically.

These successive transitions in the growth of biomass (Fig. 1 (c and d)) from the initially inactive phase to the active, inactive, and death phases correspond to those observed among the lag, exponential, stationary, and death phases. As the initial condition was chosen as the inactive phase under a condition of rich substrate availability, most of the component A molecules are arrested in a complex at this point. Therefore, at the initial stage, dissociation of the complex into component A and component B progresses, and biomass is barely synthesized, even though a sufficient and plentiful amount of substrate is available. After the cell escapes this waiting mode, catalytic reactions driven by component A progress, leading to an exponential increase in biomass. Subsequently, the external substrate is depleted, and cells experience another transition from the active to inactive phase. At this point, the biomass only decreases slowly owing to the remaining substrate and the stability of the complex. However, after the substrate is depleted and components A and B are dissociated from the complex, the biomass decreases at a much faster rate, ultimately entering the death phase.

In the active phase with exponential growth, the present model exhibits classical growth laws, namely **(i)** Monod’s growth law, and **(ii)** growth rate vs. ribosome fraction (see Fig. 6).

### Lag time dependency on starvation time *T*_stv_ and maximum growth rate *μ*_max_

In this section, we uncover the quantitative relationships among the basic quantities characterizing the transition between the active and inactive phases; i.e., lag time, starvation time, and growth rates. We demonstrate that the theoretical predictions agree well with experimentally observed relationships.

First, we compute the dependency of lag time (λ) on starvation time (*T*_stv_). Up to time *t* = 0, the model cell is set in a substrate-rich condition, 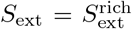, and stays at a steady state with exponential growth. Then, the external substrate is depleted to 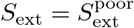 instantaneously. The cell is exposed to this starvation condition up to starvation time *t* = *T*_stv_. Subsequently, the substrate concentration *S*_ext_ instantaneously returns to 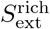. After the substrate level is recovered, it takes a certain amount of time for a cell to return to its original growth rate (Fig. S1), which is the lag time *λ* following the standard definition of lag time as the time period before the specific growth rate reaches its maximum value introduced by Penfold and Pirt[36, 37]. Given this, the dependency of λ on the starvation time *T*_stv_ can be computed.

Next, we compute the dependency of the lag time λ on *μ*_max_. We choose the steady-state solution of the cell model under 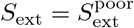 as the initial condition and compute the lag time λ under the 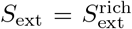 condition against different values of *μ*_max_(= *v*) (following the standard method to measure the relationship between λ and *μ*_max_[38]).

### Relationship between lag and starvation time: 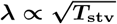

We found that λ increases in proportion to 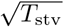, as shown in Fig. 2(a). For comparison, the experimentally observed relationship between λ and *T*_stv_ is also plotted in Fig. 2(b), using reported data [12, 18, 39] that also exhibited 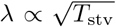 dependency. Although this empirical dependency has been previously discussed[12], its theoretical origin has thus far not been uncovered.

**FIG 2.**
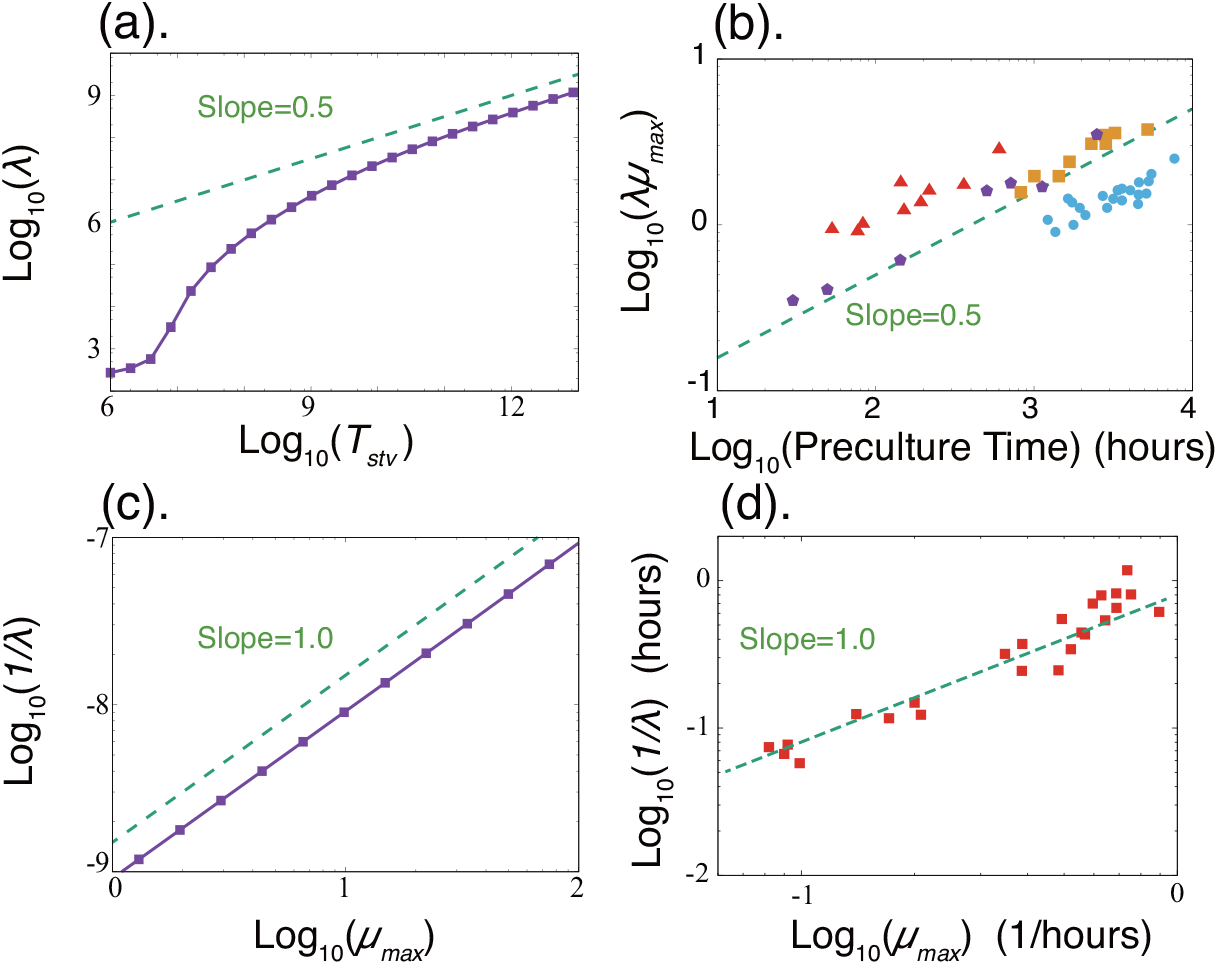
(a and b) Lag time is plotted as a function of (a) starvation time or (b) pre-incubation time. Lag time is scaled by the maximum growth rate (Inversely proportional to the shortest doubling time in the substrate-rich condition). Purple pentagons, cyan dots, and orange squares are adopted from Figures 3, 6a, and 6b of Augustin et al.[12], respectively, and the red triangles are extracted from the data in Table 1 of Pin et al.[39]. (c and d) Relationship between the lag time and maximum specific growth rate *μ*_max_. Data are adopted from Table 1 of Oscar[38]. Parameters were set as follows: 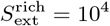, 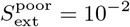, *v* = 0.1, *k_p_* = 1.0, *k_m_* = 10^−6^, *K* = 1.0, *K_t_* = 10.0, and *d_A_* = *d_B_* = *d_C_* =0 (the same parameter values as in Fig. 1 except *d_i_*s). Lag time is computed as the time needed to reach the steady state under the 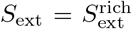 condition from an initial condition in the inactive phase. In (c), it is obtained by varying *v*(= *μ*_max_).

Indeed, the origin of 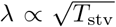 can be explained by noting the anomalous relaxation of the component B concentration, which is caused by the interaction between components A and B. A general description of this explanation is given below, and the analytic derivation is given in the *Supplementary Information*.

First, consider the time series of chemical concentrations during starvation. In this condition, cell growth is inhibited by two factors: substrate depletion and inhibition of the catalytic activity of component A. Following the decrease in uptake due to depletion of *S*_ext_, the concentration of *S* decreases, resulting in a change in the balance between A and B (hereafter we adopt the notation such that *A, B*, and *C* also denote the concentrations of the corresponding chemicals). Under the 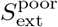 condition, the ratio of the synthesis of B to A increases. With an increase in *B, A* decreases due to the formation of a complex with B. Over time, more A becomes arrested, and the level of inactivation increases with the duration of starvation.

In this scenario, the increase of the concentration of B is slow. Considering that the complex formation reaction *A* + *B* ↔ *C* rapidly approaches its equilibrium, i.e., *k_p_AB* ~ *k_m_C*, then *A* is roughly proportional to the inverse of *B* (recall *A* + *C* = 1) if *B* is sufficiently large. Accordingly, the synthesis rate of B, given by *F_B_*(*S*)*A*, is inversely proportional to its amount, i.e.,

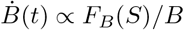

and thus

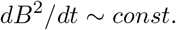

Hence, the accumulation of component B progresses with 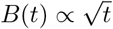. (Note that due to *S* depletion, the dilution effect is negligible.)

Next, we consider the time series for the resurrection after recovery of the external substrate. During resurrection, *A* is increased while *B* is reduced. Since component *A* is strongly inhibited after starvation, the dilution effect from cell growth is the only factor contributing to the reduction of *B*. Noting *μ* = *F_A_A* and *A* ∝ 1/*B*, the dilution effect is given by *μB* = *F_A_AB* ∝ *B*/*B* = *const*. at the early stage of resurrection. Thus, the resurrection time series of *B* is determined by the dynamics

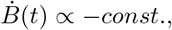

leading to the linear decrease of *B*, i.e., *B*(*t*) ~ *B*(0) − *const*. × *t*.

Let us briefly recapitulate the argument presented so far. The accumulated amount of component B is proportional to 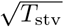, whereas during resurrection, the dilution of B progresses linearly with time, which is required for the dissociation of the complex of A and B, leading to growth recovery. By combining these two estimates, the lag time satisfies 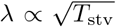.

### Relationship between the lag time and maximal growth rate: λ ∝ 1/*μ*_max_

Second, the relationship λ ∝ 1/*μ*_max_ is obtained by numerical simulation of our model, in line with experimental results [38] (Fig. 2(c and d)).

This relationship λ ∝ 1/*μ*_max_ can also be explained by the characteristics of the resurrection time series. The dilution rate of B over time is given by *μB*, as mentioned above; thus, at the early stage, *Ḃ* ~ −*μB*. In the substrate-rich condition, the substrate abundances are assumed to be saturated, so that

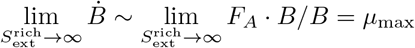

holds because lim_*S*→∞_ *F_A_*(*S*) = *μ*_max_ is satisfied. Thus, it follows that λ ∝ 1/*μ*_max_. We also obtained an analytic estimation of the lag time as

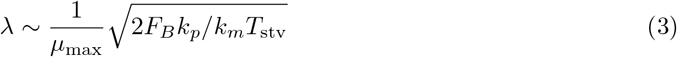

(see the *Supplementary Information* for conditions and calculation). In this form, the two relationships 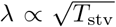 and λ ∝ 1/*μ*_max_ are integrated.

### Dependence of lag time on the starvation process

So far, we have considered the dependence of lag time on the starvation time. However, in addition to the starvation period, the starvation process itself, i.e., the speed required to reduce the external substrate, has an influence on the lag time.

For this investigation, instead of the instantaneous depletion of the external substrate, its concentration is instead gradually decreased over time in a linear manner over the span *T*_dec_, in contrast to the previous simulation procedure, which corresponds to *T*_dec_ = 0. Then, the cell is placed under the substrate-poor condition for the duration *T*_stv_ before the substrate is recovered, and the lag time λ is computed[40]

The dependence of the lag time λ on *T*_stv_ and *T*_dec_ is shown in Fig. 3(a). While λ monotonically increases against *T*_stv_ for a given *T*_dec_, it shows drastic dependence on *T*_dec_. If the external concentration of the substrate is reduced quickly (i.e., a small *T*_dec_), the lag time is rather small. However, if the decrease in the external substrate concentration is slow (i.e., a large *T*_dec_), the lag time is much longer. In addition, this transition from a short to long lag time is quite steep.

**FIG 3.**
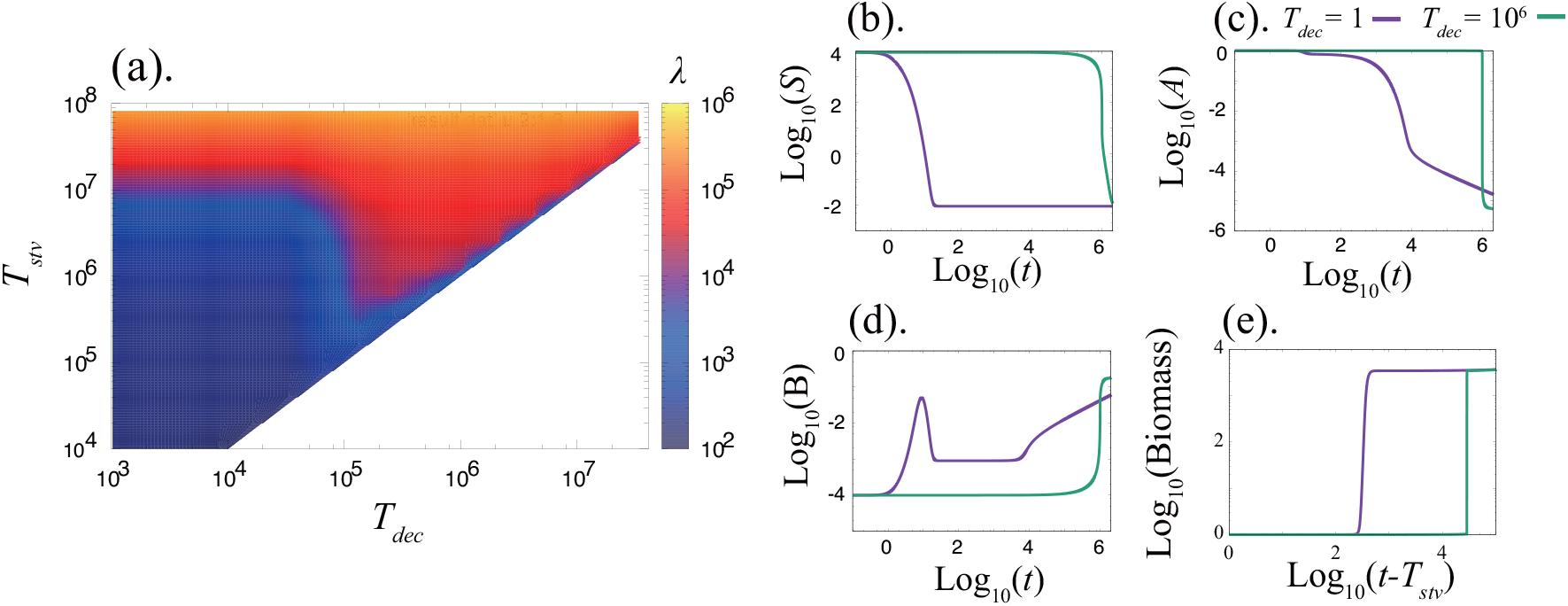
(a) Dependence of lag time λ on the time required to decrease the substrate *T*_dec_ and starvation time *T*_stv_. (b-d) Time series of starvation for different *T*_dec_ (*T*_dec_ = 10^6^ (green) and *T*_dec_ = 1.0 (purple)) values, the internal concentrations of substrate *S* (b), component A (c), and component B (d). (e) Time series of biomass during resurrection. The same parameter values as indicated in Fig.2 were adopted. The batch culture model (which is used to compute a bacterial growth curve) was adopted to compute the time series of biomass accumulation(e).Time series of *μ* is shown in Fig.S3.

This transition against the timescale of the environmental change manifests itself in the time series of chemical concentrations (see Fig. 3(b)). With rapid environmental change, *S* decreases first, whereas with slow environmental change, component A decreases first. In addition, the value of component B is different between the two cases, indicating that the speed of environmental change affects the degree of inhibition, i.e., the extent to which component A is arrested by component B to form a complex.

Now, we provide an intuitive explanation for two distinct inhibition processes. When *S*_ext_ starts to decrease, a cell is in the active phase in which A is abundant. If the environment changes sufficiently quickly, there is not enough time to synthesize the chemicals A or B, because of the lack of S, and the concentrations of chemical species are frozen near the initial state with abundant A. However, if the rate of environmental change is slower than that of the chemical reaction, the concentration of B (A) increases (decreases). Hence, A remains rich in the case of fast environmental change, whereas B is rich for a slow environmental change. In the former case, when the substrate is increased again, component A molecules are ready to work, so that the lag time is short, which can be interpreted as a kind of ”freeze-dry” process. Note that the difference in chemical concentration caused by different *T*_dec_ values is maintained for a long time because in the case of slow (fast) environmental change, chemical reactions are almost completely halted due to the decrease of *A*(*S*). Thus, the difference of lag time remains even for large *T*_stv_, as shown in Fig. 3(a).

This lag time difference can also be explained from the perspective of dynamical systems[41]. For a given *S*, the temporal evolution of *A* and *B* is given by the flow in the state space of (*A, B*). Examples of the flow are given in Fig. 4. The flow depicts (*dA*/*dt,dB*/*dt*), which determines the temporal evolution. The flow is characterized by *A* – and *B* – nullclines, which are given by the curves satisfying *dA*/*dt* = 0 and *dB*/*dt* = 0, as plotted in Fig. 4.

**FIG 4.**
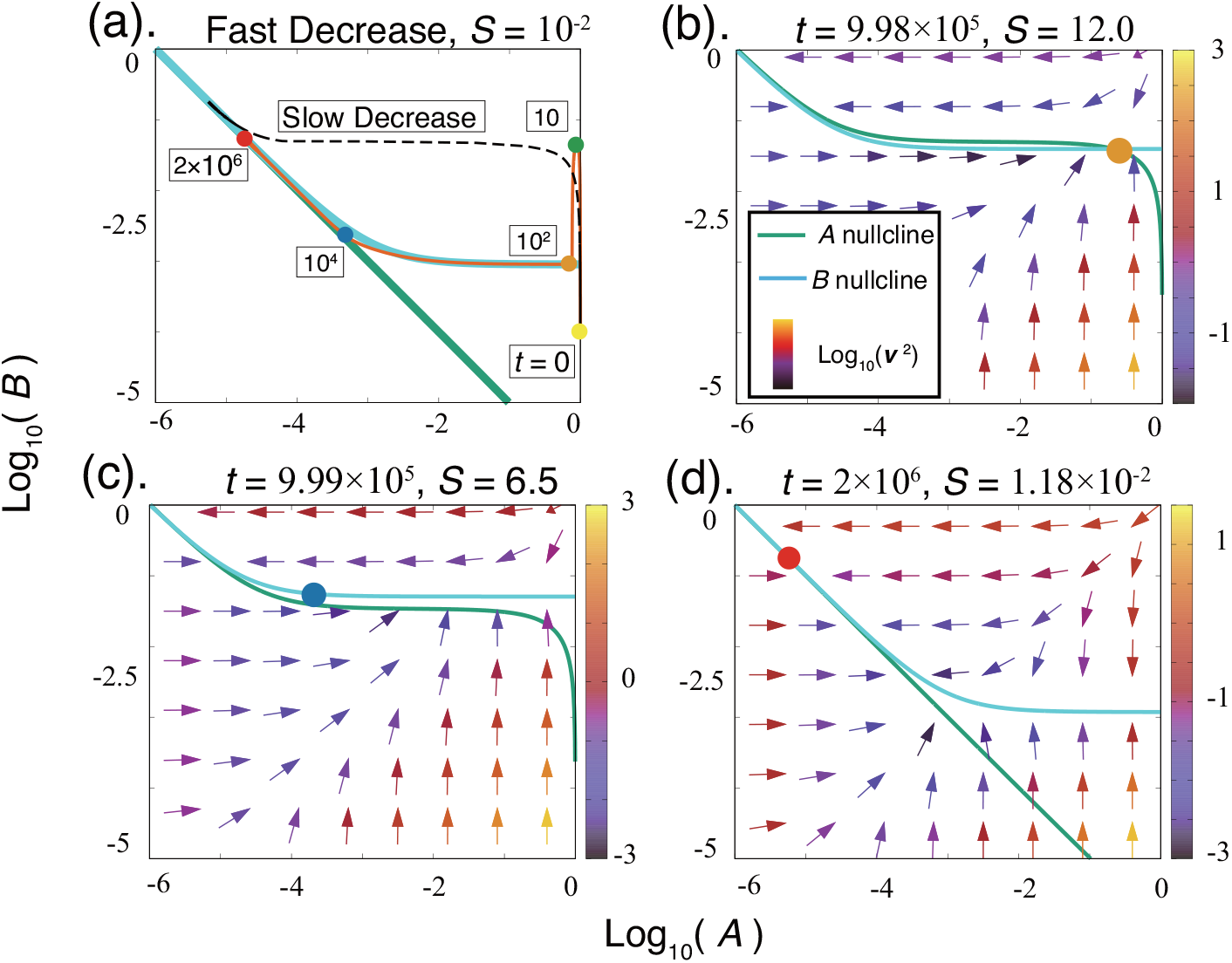
Movement of nullclines and time evolution of state variables (circles within the state space (*A, B*)). (a) The case of a fast substrate decrease (the orange line indicates the orbit and numbers in white boxes indicate the time points). The orbit of a slow substrate decrease is also plotted (black dashed line). (b–d) The case of a slow substrate decrease. Each point is the value of the state variable at the indicated time and substrate concentration. The vector field *v* = (*dA*/*dt,dB*/*dt*) is also depicted. Parameters are identical to those described in Fig. 2.

Note that at a nullcline, the temporal change of one state variable (either *A* or *B*) vanishes.

Thus, if two nullclines approach each other, then the time evolution of both concentrations *A* and *B* are slowed down, and the point where two nullclines intersect corresponds to the steady state. As shown in Fig. 4, nullclines come close together under the substrate-depleting condition, which provides a dynamical systems account of the slow process in the inactive phase discussed so far.

For a fast change (i.e., small *T*_dec_, Fig. 4(a)), *S* is quickly reduced at the point where the two nullclines come close together. First, *B* reaches the *B*-nullcline quickly. Then, the state changes along the almost coalesced nullclines where the dynamics are slowed down. Thus, it takes a long time to decrease the *A* concentration, so that at resumption of the substrate, sufficient *A* can be utilized.

In contrast, for a slow change (i.e., large *T*_dec_), the flow in (A, B) gradually changes as shown in Fig. 4(b–d). Initially, the state (A, B) stays at the substrate-rich steady state. Due to the change in substrate concentration, two nullclines moderately move and interchange their vertical locations. Since the movement of nullclines is slow, the decrease in *A* progresses before the two nullclines come close together (i.e., before the process is slowed down). The temporal evolution of *A* and *B* is slowed down only after this decrease in *A* (Fig. 4(c and d)). Hence, the difference between cases with small and large *T*_dec_ is determined according to whether the nullclines almost coalesce before or after the *A* decrease, respectively.

These analyses allow us to estimate the critical time for a substrate decrease 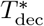 beyond the point at which λ increases dramatically. The value of a fixed point (*A*_st_, *B*_st_) depends on the substrate concentration, which drastically changes at the active-inactive transition point. If the relaxation to the fixed point is faster than the substrate decrease *T*_dec_, the system changes ‘adiabatically’ to follow the fixed point at each substrate time during the course of a “slow decrease”. The relaxation time is estimated by the smallest eigenvalue around the fixed point at the transition point. In *k_m_* → 0 limit, this eigenvalue is equal to the growth rate at the active-inactive transition point. Since it is inversely proportional to *v*, the critical time 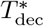 for the substrate decrease is estimated as 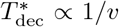. This dependence was also confirmed numerically (Fig.S4 in *Supplementary Information*).

### Distribution of lag time

So far, we have considered the average change of chemical concentrations using the rate equation of chemical reactions. However, a biochemical reaction is inherently stochastic, and thus the lag time is accordingly distributed. This distribution was computed by carrying out a stochastic simulation of chemical kinetics using the Gillespie algorithm[42].

By increasing the starvation time, two types of lag time distributions are obtained: (1) a skewed type, and (2) a skewed type with an exponential long time tail type. Each distribution type changes as follows:

(1) When the starvation time is sufficiently long, the system enters the phase with the slow accumulation of B. Here, the relaxation is anomalous, leading to a skewed type distribution. This skewed distribution is understood as follows. The number of component A molecules among cells takes on a Gaussian-like distribution just before the recovery of the external substrate concentration[43], whereas the lag time λ is proportional to *B* and thus to 1/*A*, as discussed in last section. Then, the lag time distribution λ is obtained as the transformation of 1/*A* → λ from the Gaussian distribution of component *A*. This results in a skewed distribution with a long time tail as shown in Fig. 5(a).
(2) When the starvation time is too long, the decrease in *A* comes to the stage where its molecular number reaches 0 or 1. This results in a long time tail in the distribution. This effect occurs when the number of component A molecules becomes zero due to the inhibition by component B. When the number of component A molecules becomes zero, the only reaction that can take place is a dissociation reaction (*C* → *A* + *B*). Since we assume that the time evolution of molecule numbers follows a Poisson process, the queueing time of dissociation obeys an exponential distribution Prob(queueing time = *t*) ~ *N_C_k_m_* exp(−*N_C_k_m_t*), where *N_C_* is the number of complexes formed. This exponential distribution is added to the skewed distribution, resulting in a long tail.

**FIG 5.**
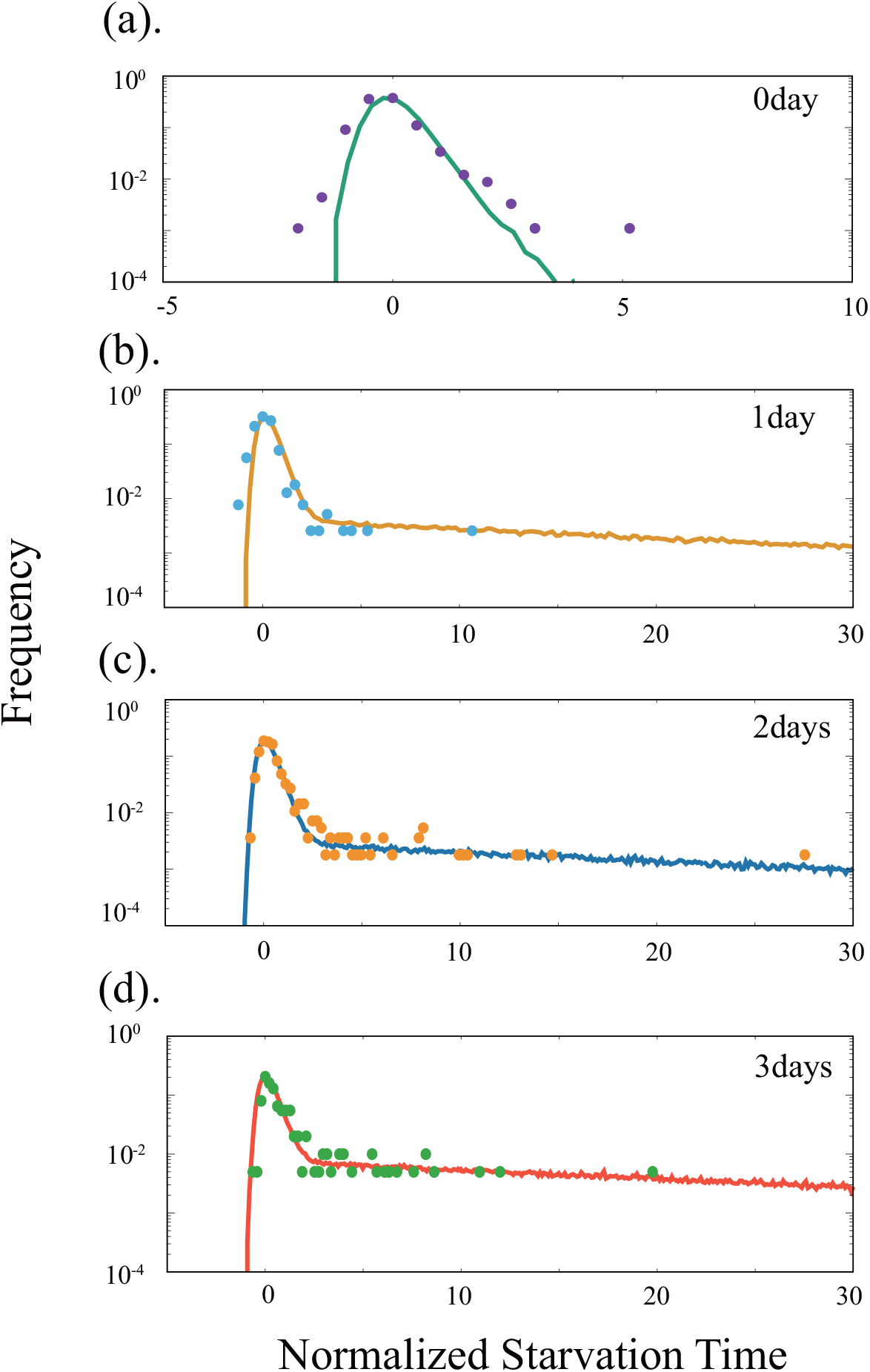
Distribution of lag time obtained by model simulation (solid line) with experimental data (lag time distribution of cultures starved for the indicated days) overlaid. The horizontal axis of each distribution was normalized by using its peak point (Peak) and the full width half maximum (FWHM) as λ → (λ − Peak)/FWHM. Experimental data were extracted from those presented in Fig. 1e of Reismann et al.[18]. Methods of stochastic simulations, the procedure used to compute Peak and FWHM, and parameter values are given in Appendix C.

The distributions of the two cases are plotted in Fig. 5, together with experimental data adopted from[18]. The skewed distribution fits the experimental observations for the 0-day starvation data, whereas the distribution including the exponential tail is a good fit to the 1-day, 2-day, and 3-day distributions.

Here, each kinetic parameter alters the critical starvation time around which the shape of the distribution starts to change; for example, a small *k_m_* makes it easier to obtain the type three distribution. However, kinetic parameters do not change the shape of the distribution directly as confirmed computationally.

The distribution of lag time was traditionally thought to follow the normal distribution[8, 44] until single-cell measurements for a long time span were carried out[18]. The preset model also generates the normal distribution of lag time if the starvation time is too short, whereas the normal distribution of lag time in earlier experiments would originate from the limitation of experimental procedures. For example, a cell that regains growth in a colony ends up dominating the colony, and thus the fluctuation of the shortest lag time governs the behavior. However, identification of the small fraction of bacteria with a long lag time is difficult owing to the limited capacity of cell tracking (as indicated in [18]).

### Remarks on the choice of parameters to fit the experimental data

Although there are several parameters in the model and the results depend on these values, the basic results on the active-inactive transition, suppression of growth, and quantitative relationships with lag time are obtained for a large parameter region. Conditions of the parameter values to obtain these main results are given in the *Supplemental Information* and are summarized in Table I. Here, an important parameter is *k_m_*, which we assumed to be the smallest among all other parameters values. This choice was made to facilitate analytic calculations, and this condition for *k_m_* can be relaxed. For example, we plotted the growth rate at the steady state in Fig.S5, indicating that the active-inactive transition occurs as long as *k_m_* < *k_p_* holds.

**TABLE I.** Predictions and Assumptions

| Result | Assumption | Condition (Prediction) |
| --- | --- | --- |
| Active-Inactive Transition Point | $k_m \sim 0$ | $1 + 2F_A(S_{\text{st}}(S_{\text{ext}}^{\text{act-inact}}))/k_p$<br>$= \sqrt{1 + 4F_B(S_{\text{st}}(S_{\text{ext}}^{\text{act-inact}}))/k_p}$ |
| Inactive-Death Transition Point | — | $F_A(S_{\text{st}}(S_{\text{ext}}^{\text{inact-death}})) = d_A$ |
| Analytic Estimation of Lag Time | $k_m \sim 0$, Dynamics of $S$ is faster than $(A, B)$ | $\lambda \sim \frac{1}{F_A(S_{\text{st}}(S_{\text{ext}}^{\text{rich}}))} \sqrt{2F_B(S_{\text{st}}(S_{\text{ext}}^{\text{poor}}))k_p/k_m T_{\text{stv}}}$ |
| Contiguity of Nullclines (Slow Relaxation) | — | $A_{A-\text{nullcline}}(B) \sim \frac{G(0,B)}{F_A(S) - G'(0,B)}$<br>$A_{B-\text{nullcline}}(B) \sim \frac{G(0,B)}{F_B(S) - F_A(S)B - G'(0,B)}$ |

Next, we estimated realistic parameter values such as the value of *v* from the literature. However, several parameter values could not be estimated directly from experimentally reported data because this would require quantitative studies at the stationary phase, which are not currently available. Thus, we estimated other parameter values by fitting Monod’s growth law [1] as well as from the reported relationship between the ribosome fraction and growth rate[4, 45, 46] (Fig.6) [47]. Since the number of parameters is greater than the minimum number required to fit the two laws in Fig. 6, the choice of parameter values is not unique. A possible set of of parameter values is listed in Table II in Appendix D.

**FIG 6.**
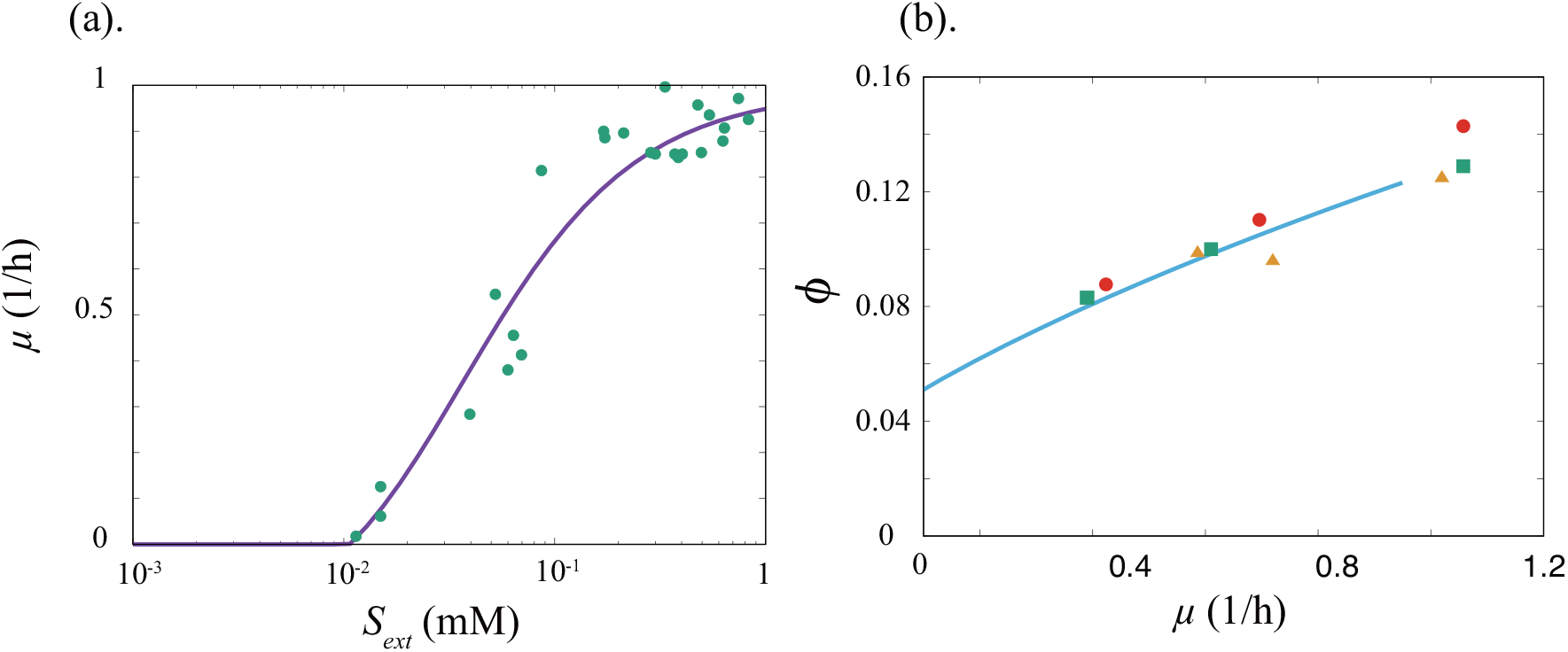
Comparison of the model results using estimated values (Table II) with experimental values. (a) The specific growth rate is plotted as a function of the external substrate (glucose) concentration. Experimental data are adopted from Monod[1]. (b)Fraction of ribosomal proteins (component A) to total proteins as a function of the specific growth rate: orange squares are from Scott et al.[4], red circles are from Bremer and Dennis[45], and green triangles are from Forchhammer et al.[46]. In (b), the theoretical curve from the model is plotted up to ~1.0, because we obtained the parameter values by fitting the *μ* − *ϕ* relation and the Monod equation with the maximum growth rate of *μ*_max_ ~ 1.0.

In fitting the two growth laws in Fig.6, we have also found that *v* is proportional to the maximum growth rate and negatively correlates to the slope of the linear relationship between ribosome fraction and growth rate, while *r* (the fraction of actively translating ribosome) decreases and *k_p_* increases the y-offset of the linear relation, respectively[48]. These relationships among parameters are consistent with those reported in[1, 4].

## DISCUSSION

Here, we developed a coarse-grained model consisting of a substrate, autocatalytic active protein (component A), non-growth-facilitating component (component B), and A-B complex, C. In the steady state, the model shows distinct phases, i.e., the active, inactive, and death phases. In addition, the temporal evolution of total biomass is consistent with the bacterial growth curve. The present model not only satisfies the already-known growth laws in the active phase but also demonstrates two relationships, 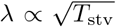 and λ ∝ 1/*μ*_max_, concerning the duration of the lag time *λ*. Although these two relationships have also been observed experimentally, their origins and underlying mechanisms had not yet been elucidated. The present model can explain these relationships based on the formation of a complex between components A and B, whose increase in the starvation condition hinders the catalytic reaction.

The above two laws are also generally derived for the inactive phase, which corresponds to the stationary phase, as long as the ratio of the synthesis of component B to that of component A is increased along with a decrease in the external substrate concentration. This condition can also be interpreted as a natural consequence of the waste-reducing (or error-correcting) process that is ubiquitous in a cell, which demands energy when assuming that component B consists of waste molecules. These laws are also derived if the waste is interpreted as a product of erroneous protein synthesis, where a proofreading mechanism to correct the error, which also requires energy, works inefficiently in a substrate-poor condition. The inhibition of growth by waste proteins is experimentally discussed by Nucifora et al. and others[26–28]. Aggregation of such waste proteins can inhibit the catalytic activity of proteins, although its role in the transition to the inactive phase remains to be elucidated. Alternatively, instead of waste proteins, we can also interpret such non-autocatalytic proteins as specific inhibitory molecules binding ribosomes such as YfiA and HPF [29–31].

For a simpler model, one could eliminate the substrate dependence of *F_B_*(*S*)/*F_A_*(*S*). Indeed, even in this simpler form, the active/inactive transition itself is observed if we tune the parameter values finely, as the decrease in substrate flow decreases the dilution, which in turn increases the fraction of complexes formed. Nevertheless, the accumulation of non-autocatalytic proteins is not facilitated with a substrate decrease, and the increase in the lag time as 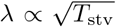 does not follow. Hence, this simpler model will not be appropriate to explain the behavior of the present cells, although it might provide relevant insight as a general mechanism for the “inactive” or “dormancy” phase in the context of protocells.

Although the cell state with exponential growth has been extensively analyzed in previous theoretical models, the transition to the phase with suppressed growth has thus far not been theoretically explained. Our model, albeit simple, provides an essential and general mechanism for this transition with consideration of the complex formation between components A and B, which can be experimentally tested.

The model here may also be relevant to study growth arrest such as stringent response[49, 50]. There, ppGpp, the effector molecule of the stringent response, is known to destabilize the open complex of all promoters causing the global reduction of macromolecular synthesis, playing the similar role as the component B in the present paper[51–54]. Additionally, rpoS, sigma factor of stationary-phase genes, lies downstream of ppGpp[55], and it is reported that the mutant lacking ppGpp (which might correspond to inhibition of the component B in our model) shows a physiological state reminiscent of exponentially growing bacteria even under starvation [56].

Moreover, the model predicts that the lag time differs depending on the rate of external depletion of the substrate, which can also be examined experimentally. Recently, the bimodal distribution of growth resumption time from the stationary phase was reported in a batch culture experiment[19]. The heterogeneous depletion of a substrate due to the spatial structure of a bacterial colony is thought to be a potent cause of this bimodality, and progress toward gaining a deeper understanding of this concept is underway. Since the present model shows different lag times for different rates of environmental change, it can provide a possible scenario for helping to explain this bimodality.

## ACKNOWLEDGEMENT

The authors would like to thank S. Krishna, S. Semsey, N. Mitarai, A. Kamimura, N. Saito, and T. S. Hatakeyama for useful discussions; and I. L. Reisman, N. Balaban, and J.C. Augustin for providing data. This research is partially supported by the Platform for Dynamic Approaches to Living System from the Japan Agency for Medical Research and Development (AMED), Grant-in-Aid for Scientific Research (S) (15H05746 from the Japan Society for the Promotion of Science [JSPS]), and JSPS grant 16J10031.

## APPENDIX A: MODEL WITHOUT INTERACTION BETWEEN THE TWO COMPONENTS

To clarify the necessity of the interaction between the two components to obtain the main results, we remove the complex formation between A and B (by setting *k_p_* and *k_m_* to be zero).

Then, the A-B complex is eliminated, and our model is given as

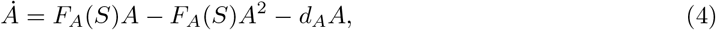

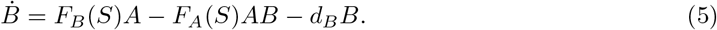

(We assume that the internal concentration of the substrate is equal to that of the external concentration of the substrate, and ignore the substrate dynamics.) The steady solution is

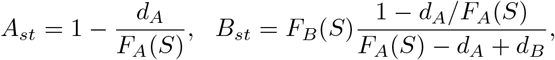

and the steady growth rate is given as *μ_st_* = *F_A_*(*S*)*A_st_* = *F_A_*(*S*) − *d_A_*. Therefore, the present model without an interaction between components A and B exhibits only the active-death transition at *S*\*, satisfying *F_A_*(*S*\*) = *d_A_*.

In addition, the dynamics of the system are calculated as

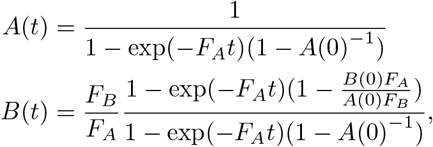

where we neglect *d_i_*, as in the main text. Therefore, if the model cell Eq. (5) restarts growth in a high *S* (*S_rich_*) value environment after exposure to the starvation condition (low *S* value), *A*(*t*) and *B*(*t*) exponentially converge to the substrate-rich steady state. Hence the time for growth recovery *T*_rec_ is quite short, which is calculated

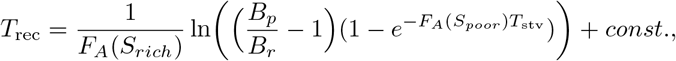

as a function of starvation time *T*_stv_. Here, *B_p_* and *B_r_* are the steady concentrations of component *B* under the substrate-poor and substrate-rich environment, respectively. Obviously this relationship is far from the relationship between lag and startvation time.

## APPENDIX B: REDUCTION OF THE KINETIC PROOFREADING MODEL

In the main text, the concrete forms of *F_A_* and *F_B_* were predetermined by assuming the characteristic 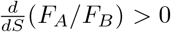, which is essential for the active-inactive transition. In this section, we show that this characteristic is derived from a simple polymer elongation model with a kinetic proofreading scheme[33] by assigning a correct polymer as A and an erroneous one as B. Indeed, 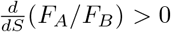 originates from an error in the synthesis of component A that consequently inhibits the synthetic reactions.

Polymer elongation is essential to synthesize macromolecules. It is well known that ribosomes elongate a polypeptide chain following receipt of the information from messenger RNA. However, since the transfer RNA (tRNA) discrimination by a ribosome is not perfect, there is always a certain probability for mistranslation (i.e., the wrong choice of tRNA). Kinetic proofreading is one of the possible error-correction mechanisms in such a polymerization system, which demands energy. We derive that the synthesis ratio of mistranslated proteins to a “correct” protein increases under the substrate-depleting condition.

For the polymerization reaction, we introduce two monomers, “correct” and “wrong” monomers, as simplified from real amino acids. In reality, there are 20 amino acids and one tRNA that specifies one amino acid, i.e., one correct and 19 wrong monomers with a certain affinity lower than that of the correct monomer.

In the model, a polymer is elongated up to the length *L* with the aid of the catalytic activity of the “correct” protein, i.e., the ribosome. The matured polymer with length *L* is spontaneously folded into a protein; the proteins consisting of only correct monomers are correct proteins with catalytic activity, whereas those with other monomer sequences turn into mistranslated proteins. The elongation process progresses under a kinetic proofreading mechanism (Fig. 7). As in the original model, mistranslated proteins inhibit the correct protein’s catalytic activity by forming a complex with it, while the growth is facilitated by the activity of correct proteins.

**FIG 7.**
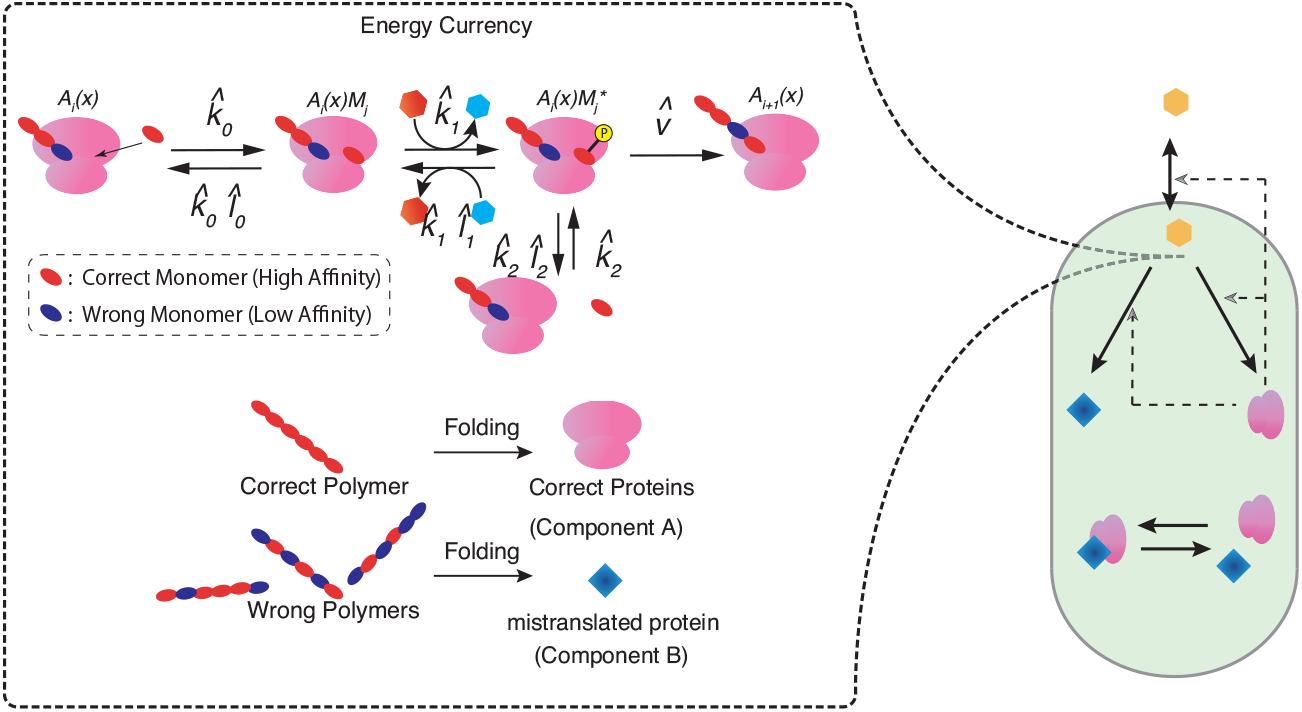
Schematic representation of a polymer elongation system with kinetic proofreading. The reactions other than the synthesis part (*F_A_*(*S*)*A* and *F_B_*(*S*)*A*) are identical to those of the original model (2).

The dynamics of the polymer elongation part are given by

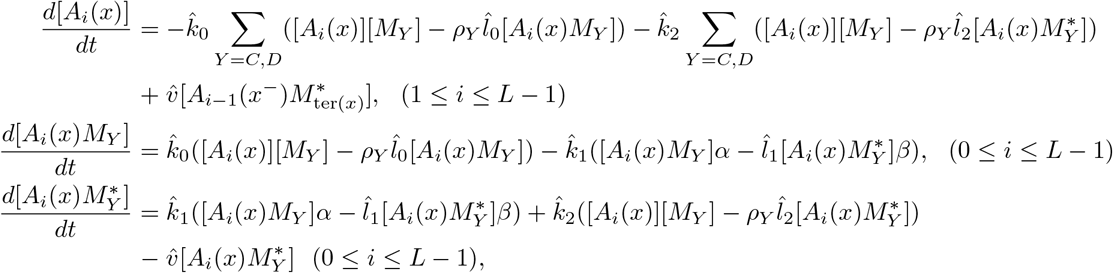

where [*M_C_*] and [*M_D_*] denote the concentrations of correct and wrong monomers, respectively. [*A_i_*(*x*)], [*A_i_*(*x*)*M_Y_*], and 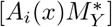 represent the concentration of a complex of correct proteins and a polymer with length i, a correct protein-polymer-monomer complex, and an activated correct protein-polymer-monomer complex, respectively, where *x* denotes a monomer sequence such as *CCDC* ⋯, with *C* and *D* indicating the correct and wrong monomer, respectively. ter(*x*) and *x*^−^ indicate the last monomer (*C* or *D*) of a monomer sequence *x* and the partial monomer sequence of *x* from which the last monomer (i.e., ter(*x*)) has been removed, respectively. Here, [*A*_0_] denotes the concentration of the correct protein. 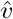 and 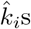 are the rate constants of the chemical reactions, and the 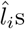 are the Boltzman factors of each chemical reaction. We assume that dissociation of the matured polymer from correct proteins and polymer folding into proteins take place instantaneously. *α* and *β* are the concentration energy currencies, for example, GTP and GDP, respectively. *ρ_i_* reflects the difference in affinity between the wrong monomer (D) and the correct monomer (C) (we set *ρC* as unity).

At the steady state, the synthesis rates of correct and mistranslated proteins, 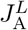 and 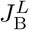, are given by

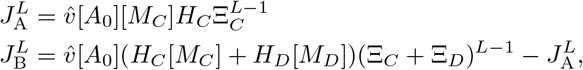

where functions Ξ_*i*_ and *H_i_* are given by

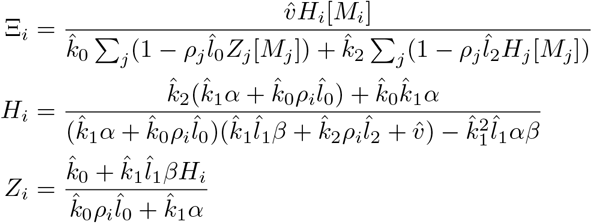

Ξ_0_ and Ξ_1_ denote the rate of polymer elongation with the wrong and correct monomers, respectively.

Now, we set the functional form of *α* and monomer concentrations [*M_C_*] and [*M_D_*] to obtain the concrete values of 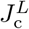 and 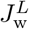. It is natural to assume that *α* and [*M_i_*] are increasing functions of the internal substrate concentration [*S*]. Here, we adopt a Michaelis-Menten’s type form *α* = [*S*]/(*K_a_* + [*S*]), *β* = *K_a_*/(*K_a_* + [*S*]), and [*M_C_*] = [*M_D_*] = [*M*]_max_[*S*]/(*K_S_* + [*S*]).

Although 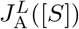 and 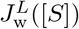 do not completely agree with the form we adopted for *F_A_*(*S*)*A* and *F_B_*(*S*)*A* in the original model, the conditions discussed in Section 2 of the Supplementary Information are nevertheless satisfied, as shown in Fig. 8. In particular, 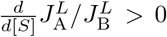 holds. Indeed, using this model, we obtained the same active, inactive, and death phases, as well as the same growth curve and other quantitative laws. As an example, Fig. 9 shows the steady growth rate as a function of the external substrate concentration [*S*]_ext_. Furthermore, for any *L*, the same behaviors are obtained, as 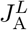 and 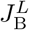 satisfy the condition outlined in Section 2 of the Supplementary Information. It is also confirmed that the ratio of 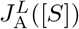 to 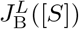 increases as [*S*] increases for any *L*.

**FIG 8.**
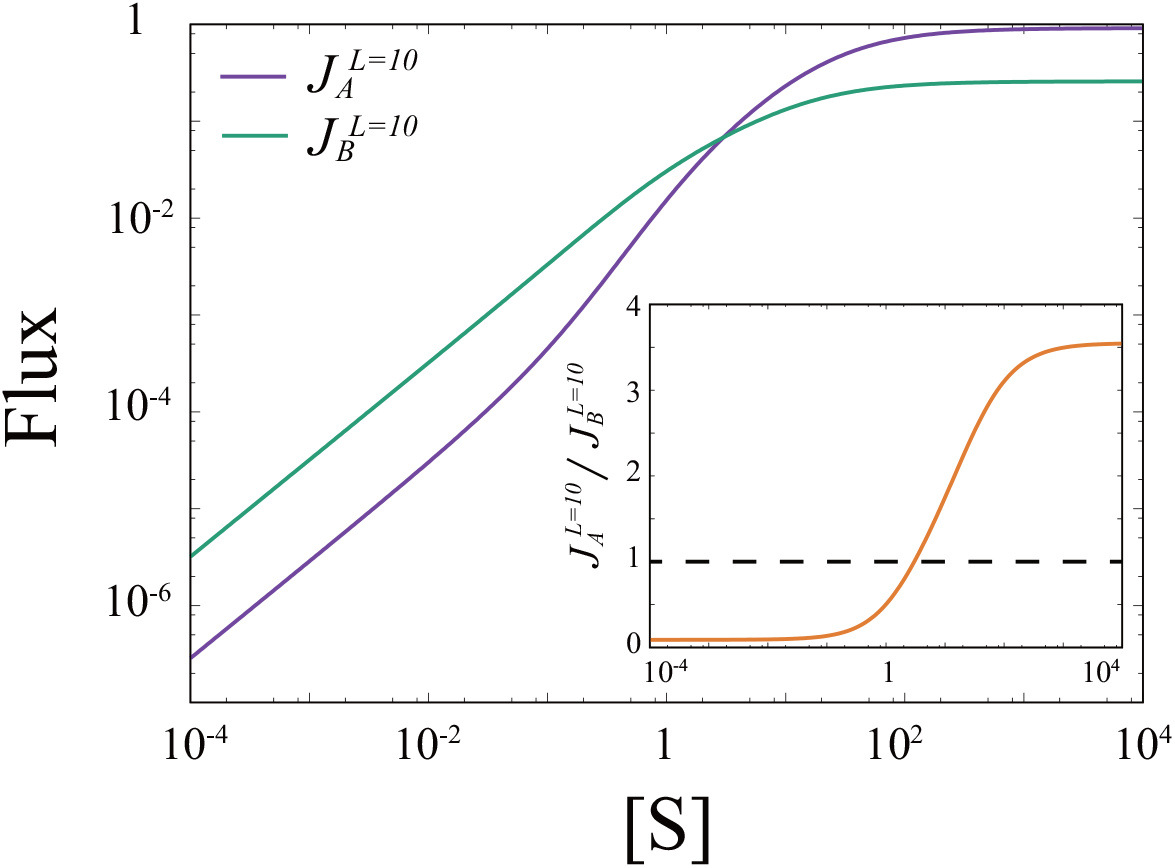
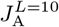 and 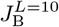 are plotted against the substrate concentration [*S*]. The ratio of 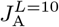 to 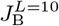 is also plotted in the inset of the figure. Parameters for the polymer elongation part are set to be 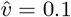, *ρC* = 1.0, *ρD* = 10.0, 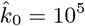, 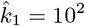, 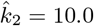, 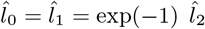, 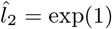, *K_a_* = 10.0, *K_S_* = 1.0, [*M*]_max_ = 1.0, and [*A*_0_] = 1.0.

**FIG 9.**
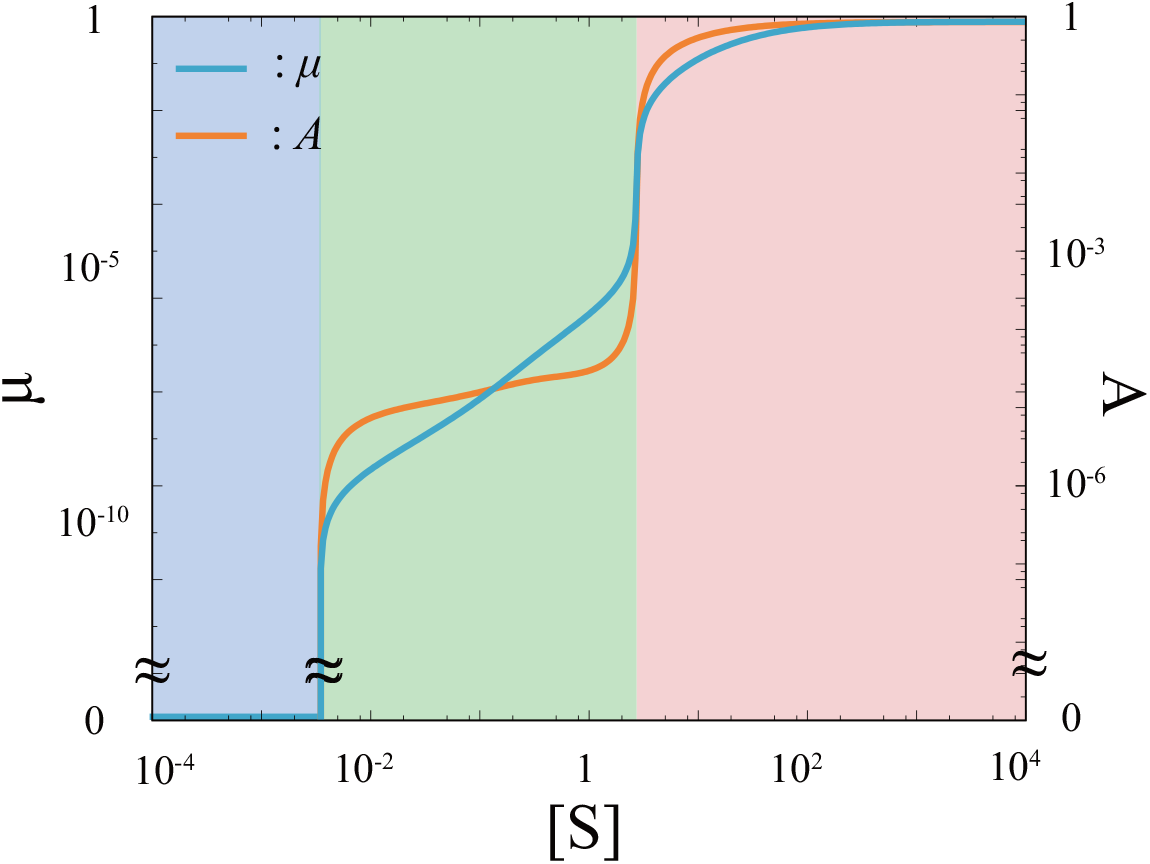
Steady growth rate of the model with polymerization and kinetic proofreading. 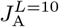 and 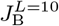 are adopted for the synthetic reaction rate of components A and B. Parameters for 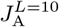 and 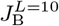 are identical to those in Fig. 8, and others are set to be the same.

## APPENDIX C: DETAILS OF MODELS AND SIMULATION PROCEDURES

To obtain the growth curve shown in Fig. 1(c) and (d) in the main text, we added the dynamics of the substrates in the external environment, as well as cell volume growth. By representing the dynamics according to the amounts of chemicals rather than their concentrations, the model is given by

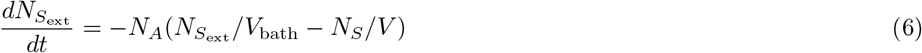

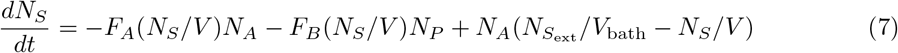

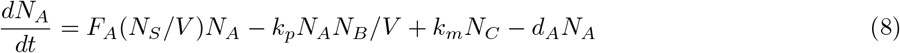

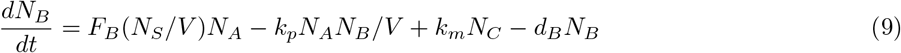

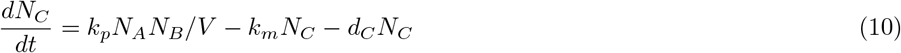

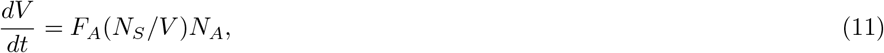

where *N*_*S*_ext__ is the amount of substrate in the external environment at volume *V*_bath_, and *N_S_, N_A_, N_B_*, and *N_C_* are the amounts of each chemical within the cell at volume *V*(*t*), respectively. *V*(*t*) is the volume of a cell. The dilution effect is introduced by dividing the amount of each chemical by *V*(*t*). *S*_ext_ is the total amount of the external substrate contained in the culture system with volume *V*_bath_ (set to be unity). For all other parameters, the same values as shown in Fig. 1 were adopted.

To obtain the lag time distribution, we performed a stochastic simulation. We computed the model equation according to the volume change

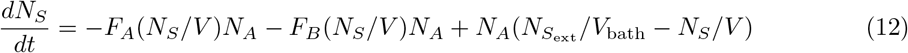

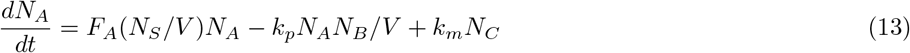

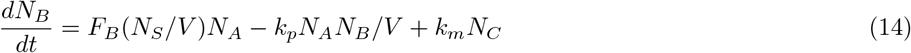

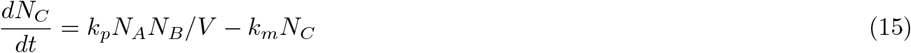

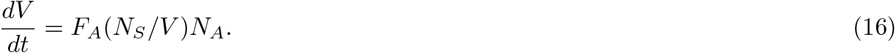

Here, we introduced cell division and simulated the dynamics of only one daughter cell (to reduce the simulation time). When the cell volume *V* reaches the division volume *V*_div_, *V* halves and chemicals are distributed to two doughter cells in equal probability. After computing these equations for a sufficiently long time under the 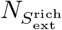 condition, *N*_*S*_ext__ suddenly changed to 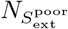, and was then set at this value over the starvation period *T*_stv_. Then, *N*_*S*_ext__ returned to the original value 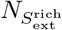. The lag time λ is computed as the time needed to double the volume from *V*_0_; i.e., the volume at which *S*_ext_ recovers. The numerical results indicated that the absolute value of the correlation coefficient between *V*_0_ and λ is small. Here, the difference in *V*_0_ in cells does not affect the distribution of the lag time. Stochastic simulation was carried out using the Gillespie algorithm. Parameter values were set to be *V*_div_ = 2 × 10^3^, *V*_bath_ = 1.0. 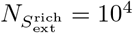, 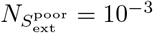, and the others were the same as those described in Fig. 2. The length of starvation time *T*_stv_ was set to be 5 x 10^4^,10^6^, 2 x 10^6^, and 10^7^ for Fig. 5 (a), (b), (c), and (d), respectively.

From the lag time distribution obtained by numerical simulation, we could compute the peak and FWHM values directly. Since the experimental data did not include a sufficient amount of samples, we applied a smoothing filter to determine the FWHM, while the peak point was determined directly.

## APPENDIX D: ESTIMATED PARAMETER VALUES

**TABLE II.** Estimated parameter values

| Symbol | Meaning | Estimated Value (Unit) | Reference |
| --- | --- | --- | --- |
| $f_0$ | Stoichiometry of glucose and amino acids (measured by carbon, average) | 1.1(-) | [57] |
| $f_1$ | Stoichiometry of amino acids and average proteins | 209(-) | [58] |
| $f_2$ | Stoichiometry of amino acids and ribosomal protein | 7336(-) | [4] |
| $v$ | synthesis rate of protein per ribosome<br>$20(\text{a.a/sec})/209(\text{a.a}) \times 3600(\text{sec})$ | 345 (1/hour) | [4, 58] |
| $m$ | volume growth per synthesis of growth factor<br>$1.0\mu\text{m}^3$ (E. coli volume) divided by<br>$5 \times 10^5$ (# of proteins) | $1.2 \times 10^{-2}$ (1/mM) | [58] |
| $r$ | Fraction of actively translating ribosomes | 0.8 (-) | [45] |
| $\psi$ | Fraction of ribosomal proteins to component A | $3.6 \times 10^{-3}(-)$ | Fitting |
| $D$ | Speed of phenomenological catabolism | $10^5(1/\text{mM}/\text{hour})$ | Fitting |
| $K$ | Saturation constant of $F_A + F_B$ | $1.8 \times 10^{-2}(\text{mM})$ | Fitting |
| $K_t$ | Phenomenological constant changing A/B balance | $2.3 \times 10^{-2}(\text{mM})$ | Fitting |
| $k_p$ | Rate of association between A and B | $10^{-3}(1/\text{mM}/\text{hour})$ | Fitting |
| $k_m$ | Rate of dissociation between A and B | $10^{-6}(1/\text{hour})$ | Fitting |

